# Characterization of LINE-1 transposons in a human genome at allelic resolution

**DOI:** 10.1101/594200

**Authors:** Lei Yang, Genevieve A. Metzger, Richard N. McLaughlin

**Affiliations:** Pacific Northwest Research Institute, Seattle, WA 98122, USA

## Abstract

The activity of the retrotransposon LINE-1 has created a substantial portion of the human genome. Most of this sequence comprises fractured and debilitated LINE-1s. An accurate approximation of the number, location, and sequence of the LINE-1 elements present in any single genome has proven elusive due to the difficulty of assembling and phasing the repetitive and polymorphic regions of the human genome. Through an in-depth analysis of publicly-available, deep, long-read assemblies of nearly homozygous human genomes, we defined the location and sequence of all intact LINE-1s in these assemblies. We found 148 and 142 intact LINE-1s in two nearly homozygous assemblies. A combination of these assemblies suggests a diploid human genome contains at least 50% more intact LINE-1s than previous estimates – in this case, 290 intact LINE-1s at 194 loci. We think this is the best approximation, to date, of the number of intact LINE-1s in a single diploid human genome. In addition to counting intact LINE-1 elements, we resolved the sequence of each element, including some LINE-1 elements in unassembled, presumably centromeric regions of the genome. A comparison of the intact LINE-1s in each assembly shows the specific pattern of variation between these genomes, including LINE-1s that remain intact in only one genome, allelic variation in shared intact LINE-1s, and LINE-1s that are unique (presumably young) insertions in only one genome. We found that many old elements (> 6 million years old) remain intact, and comparison of the young and intact LINE-1s across assemblies reinforces the notion that only a small portion of all LINE-1 sequences that may be intact in the genomes of the human population has been uncovered. This dataset provides the first nearly comprehensive estimate of LINE-1 diversity within an individual, an important dataset in the quest to understand the functional consequences of sequence variation in LINE-1 and the complete set of LINE-1s in the human population.

## Introduction

The replication of transposable elements has created much of the sequence in large genomes (Sotero-Caio et al. 2017; Canapa et al. 2015). Most of this sequence contains only the degraded versions of once active transposons. However, activity of transposable elements underlies a variety of human diseases, driven by both the disruptive nature of insertions and the immune sensing of replicating elements (Hancks and Kazazian 2012; Crow 2010). It is also clear that activity of transposons gives rise to variation amongst human genomes (Auton et al. 2015; Beck et al. 2011). Further, the permanence of transposon sequences in genomes has resulted in the co-option of these elements to support host functions through their DNA, RNA, and protein products (Kapusta et al. 2013; Chow et al. 2010; Bejerano et al. 2006; Finnerty et al. 2002; Jacques et al. 2013). As such, it is crucially important to understand with precise detail the location, sequence, and variation of transposable elements in the human genome.

In the human genome, only one class of transposons is measurably active and autonomous, encoding all the parts necessary to replicate in a host cell. These Long Interspersed Element-1s (LINE-1s or L1s) copy themselves within the host genome via reverse transcription– a process termed ‘retrotransposition’. Consequently, millions of LINE-1 copies comprise at least 17% of the contemporary human genome (Lander et al. 2001; de Koning et al. 2011; Smit et al.). The oldest detectable LINE-1 in the human genome likely predate the common ancestor of placental mammals (Smit et al. 1995); elements from the youngest lineage of human LINE-1 actively retrotranspose (Moran et al. 1996; Brouha et al. 2003) and are polymorphic in the population (Stewart et al. 2011; Beck et al. 2010; Huang et al. 2010; Iskow et al. 2010; Ewing and Kazazian 2010). Yet, the complete scope and scale of the role of LINE-1 in human genetic variation and biology is a topic of intense investigation. Previous landmark efforts to determine the number and sequence of full-length and/or putatively retrotransposition-competent LINE-1s found 89 intact LINE-1s and about half of these had measurable *in vitro* retrotransposition activity. The authors’ extrapolation to a diploid genome estimated that the human genome contains 80-100 retrotransposition-competent LINE-1s (Brouha et al. 2003). Here, intact LINE-1s were defined by their similarity to a reference sequence L1.3 (GenBank accession number: L19088) (Dombroski et al. 1993) and the preservation of two open reading frames. Of note, this study used the only resource available at the time - an early, relatively low-quality assembly of the human genome, depleted of high-quality assembly in repeat regions (human genome working draft (HGWD), December 2001 freeze).

In addition to extracting LINE-1 sequences from genome assemblies, numerous targeted approaches have provided important advancement of our understanding of LINE-1 variation amongst individuals. Individual genomes vary in their LINE-1 content at two levels: the presence/absence state, also known as ‘insertion polymorphisms’, and the allelic differences of homozygous LINE-1 insertions. Targeted, short read-based resequencing approaches to map LINE-1 insertions relative to a reference and junction-fragment sequencing have demonstrated that LINE-1s show dramatic insertional polymorphism in the human population (Tang et al. 2017; Iskow et al. 2010; Sudmant et al. 2015; Xing et al. 2009). Other groups have used whole genome BAC libraries to find and sequence LINE-1 insertions that are polymorphic relative to the human reference assembly (Kidd et al. 2010; Beck et al. 2010). Despite the importance of these approaches, they are not designed to describe the location and full-length sequences at allelic resolution of all LINE-1 elements of individual genomes.

Further, assembly and resequencing-based approaches have been plagued by a fundamental complication of sequencing diploid genomes: phasing the two alleles of each stretch of genomic DNA. This is particularly problematic if there exists significant heterozygosity at repeat sequences in the genome. While the extent of allelic variation remains unclear, case studies demonstrate that allelic variation exists at LINE-1 insertions, and this variation can have functional consequences (Lutz et al. 2003).

Due to a reliance on short reads and the unphased nature of most genome assemblies, we do not yet have a reliable description of the extent of intactness amongst all LINE-1s within individual human genomes (Figure 1). Only with recent ambitious efforts to deeply sequence nearly homozygous human cell lines with long read technologies have we gained the ability to comprehensively catalog the LINE-1 sequences in individual haploid human genomes at allelic resolution (Steinberg et al. 2014; Chaisson et al. 2015; Huddleston et al. 2017). These assemblies overcome previous limitations by using 1) reads which are, on average, longer than the 6,000bp repeat unit length of LINE-1; 2) source genomes which are homozygous, eliminating the need to phase alleles of LINE-1 repeats. In this paper, we describe our process for identifying intact LINE-1s from these assemblies and characterizing the variation that exists between two assemblies. We posit that our approach provides a nearly comprehensive catalog of these sequences which enables us to extrapolate the state of LINE-1 variation within and between genomes.

**Figure 1.**
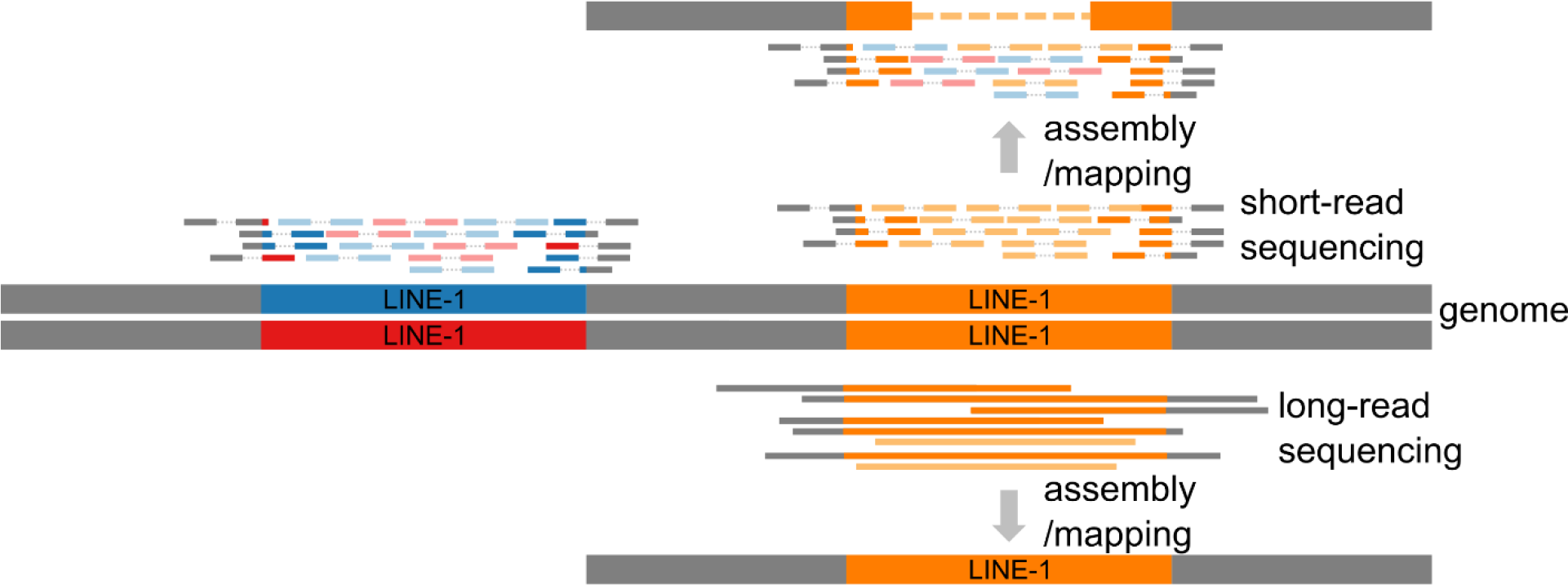
Long-read sequencing of homozygous genomes allows identification of the location and sequence of LINE-1 alleles in individual genomes. A cartoon model mapping or assembling LINE-1 sequences is shown based on sequencing with short reads (top) or long reads (bottom) at a heterozygous (blue and red) and a homozygous (orange) locus of the genome. Short reads are more likely to be completely embedded within a LINE-1 (sequencing reads with light hue), creating gaps in the assembly or ambiguous mapping (top) and only resolving the junction of insertions (sequencing reads marked by solid color and gray bars). Reads longer than LINE-1 (~6,000 bp) used to assemble nearly homozygous genomes resolve the sequence of LINE-1 alleles in the genome (bottom).

## Results

### Long read sequencing of homozygous genomes reveals the location and sequence of intact LINE-1s

The CHM1 (complete hydatidiform mole 1) assembly (Steinberg et al. 2014) represents the nearly homozygous genome (<0.75% heterozygosity) (Fan et al. 2001) of a human hydatidiform mole cell line derived from a European individual and deeply sequenced with PacBio reads (54x). Extensive characterization of this assembly has annotated the location and type of structural variants, including LINE-1s (Huddleston et al. 2017; Chaisson et al. 2015; Steinberg et al. 2014). Extending these analyses to broadly investigate the sequence of LINE-1s in this genome, we identified all LINE-1 sequences by compiling RepeatMasker (Smit et al.) annotations of the assembly and filtering for sequences masked as LINE-1. We found 919,967 sequences annotated as LINE-1, comprising ~504 Mbp of total LINE-1 sequence. This corresponds to ~16.82% of the total genome assembly, consistent with previous estimations (Chaisson et al. 2015; Lander et al. 2001).

As expected, most LINE-1 sequences of CHM1 were short fragments less than 500bp (Figure 2A). Many of the longer fragments were 5’ truncated LINE-1s, a pattern expected from the often abortive 3’-5’ target-primed reverse transcription of LINE-1s (Szak et al. 2002). We also observed internal fragments and 3’ truncated pieces. Another smaller peak was evident in a histogram of sequence lengths around 6,000 bp, the length of full-length LINE-1 sequences (Figure 2A, 2B). Within this full-length peak, we observed two peaks at around 6,100 bp and another smaller peak at around 6,400 bp. The sequences in each of these peaks differ in a 129bp deletion in their 5’UTR (the 6100 bp peaks) or a completely different 5’ UTR (the 6400 bp peak) (Jacobs et al. 2014; Khan et al. 2006).

**Figure 2.**
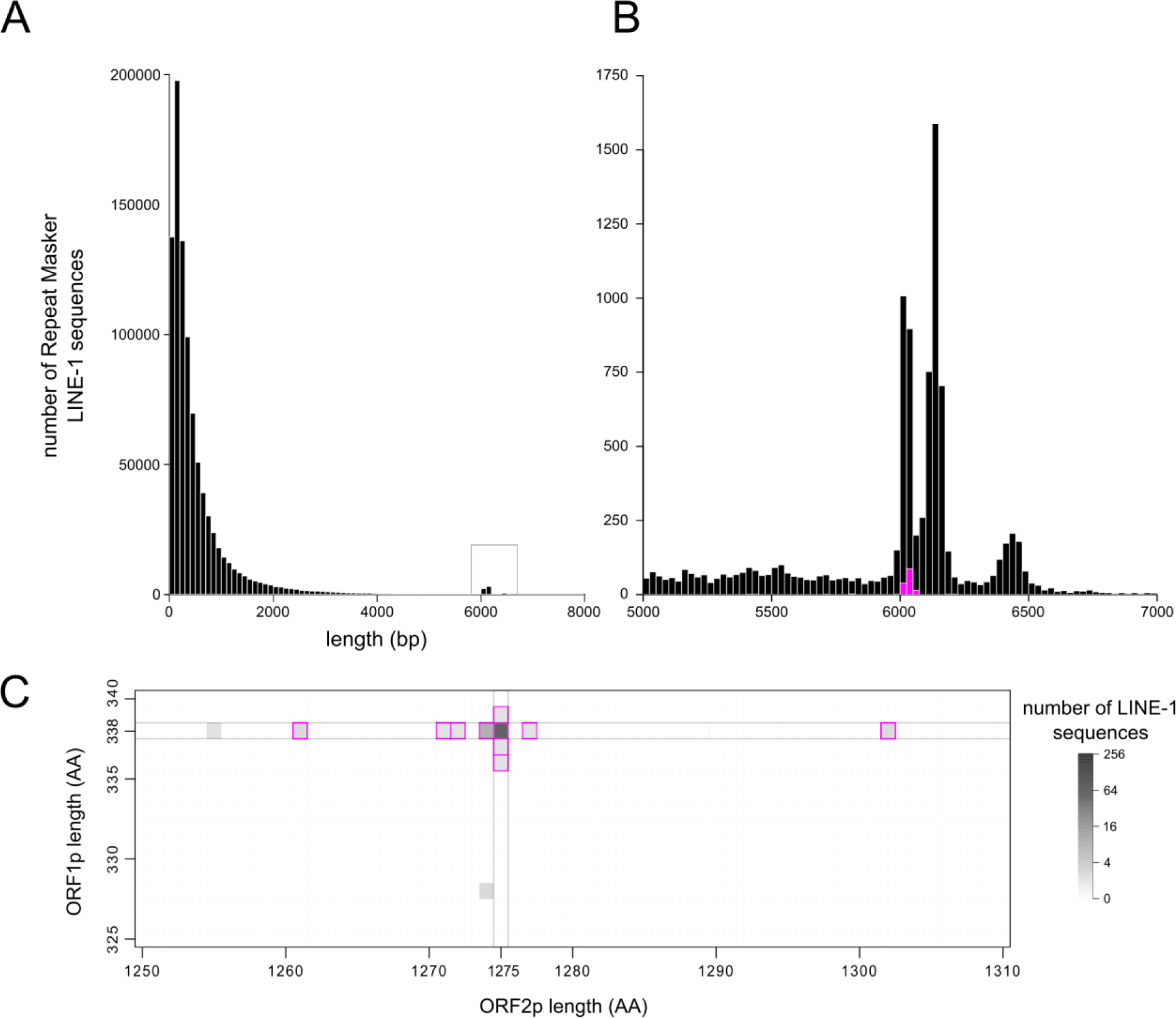
Identification of intact LINE-1s in CHM1. **A.** The distribution of lengths of LINE-1-masked sequences shows most sequences are less than 1,000 bp. An additional peak around 6,000 bp contains the full-length LINE-1 copies. **B.** A zoomed-in view of the boxed region of the complete length histogram. Intact LINE-1 sequences are highlighted in magenta. **C.** A heatmap of the number of sequences with the indicated translated ORF1 and ORF2 lengths. Within the full-length set of LINE-1 sequences, only 148 sequences (magenta boxes) encode putative ORFs that align along the entire length of the ORF1p and ORF2p sequences of a reference element (L1_RP_).

A full-length, intact LINE-1 sequence contains two open reading frames (ORFs) which encode proteins (ORF1p and ORF2p) required for replication (Moran et al. 1996). To identify full-length copies with intact open reading frames, we analyzed the lengths of the longest translated ORFs present in each identified LINE-1 sequence that aligned to sequences of ORF1p and ORF2p from a reference LINE-1, L1_RP_ (GenBank accession number AF148856) (Kimberland et al. 1999) (Figure S1). Many sequences encode ORFs of exactly the same length as L1_RP_, 338 codons for ORF1 and 1,275 codons for ORF2 (Figure 2C). These sequences align along the entire length of the amino acid sequence of the reference, from start to stop codon without terminal deletions or extensions. In addition, we included several LINE-1s that contain shorter or longer ORFs which still align to the full-length of the amino acid sequence of the ORF1p or ORF2p of L1_RP_. As such, we define intact LINE-1s as sequences greater than 5,000 bp (9,548 LINE-1s found in CHM1 using this simple length cutoff) with two intact open reading frames that align, when translated, to the full length of L1_RP_ ORF1p (338 amino acids) and ORF2p (1,275 amino acids). With this definition of an ‘intact’ LINE-1, we conclude that the CHM1 haploid genome contains 148 intact and potentially-replication-competent LINE-1s.

A genome assembly from a second human hydatidiform mole cell line has been released and similarly analyzed for variation in segmental duplications (Huddleston et al. 2017; Schneider et al. 2017). This assembly was also built with deep coverage PacBio reads (52x), and has been extensively analyzed for structural variants, though like CHM1, no one has specifically analyzed the sequence of LINE-1s in this second homozygous genome. This sample is of unknown ethnic origin, but clusters with CHM1 and other European genomes. Using our pipeline built for CHM1, we identified intact LINE-1s in CHM13. The distribution of LINE-1 sequence lengths in CHM13 was similar to CHM1, and we found 142 intact and putatively-replication-competent LINE-1s in this assembly (Figure S2).

### Comparison of CHM1 and CHM13 to other genome assemblies

Next, we compared the intact LINE-1s in the ostensibly haploid CHM genomes to the intact LINE-1s described based on a variety of sequencing, assembly, and discovery strategies. Unsurprisingly, the initial draft release of the human reference genome (HGWD) (Lander et al. 2001), the version used to make an initial estimate of the number of LINE-1s in a human genome (Brouha et al. 2003), contains ~60% as many intact LINE-1s as the CHMs (Table 1, row 5-8). The number of intact LINE-1s grows as the reference genome becomes more complete, and the most recent major release of the reference genome (GRCh38 2017, Table 1, row 8) and CHM1 contain a comparable number of intact LINE-1s. However, for the reference genome this number represents some combination of the LINE-1s found in the genome sequences of over 50 individuals (a ‘mosaic haploid’ assembly). Assemblies of individual genomes sequenced with long reads have also been published and carefully analyzed for structural variants (Audano et al. 2019). A sampling of these heterozygous diploid-based assemblies (Table 1, row 3-4) finds approximately 50% as many intact LINE-1s as CHM1/13 using our BLAST-based approach to identify intact LINE-1s.

**Table 1:** Numbers of intact LINE-1 elements from available human genomes. The number of intact LINE-1s was counted for each genome using the methodology we describe here with the exception of HGWD, which is the number of intact LINE-1s Brouha et. al reported in their 2003 paper.

| Assembly | # Intact L1 | # individuals | Seq method | Notes |
| --- | --- | --- | --- | --- |
| CHM1 | 148 | 1 | PacBio | Diploid, nearly homozygous |
| CHM13 | 142 | 1 | PacBio | Diploid, nearly homozygous |
| NA19240 | 69 | 1 | PacBio | Diploid, heterozygous |
| NA19434 | 84 | 1 | PacBio | Diploid, heterozygous |
| HGWD (2001) | 90 | - | mixed | Phased mosaic diploid |
| GRCh37 (2009) | 104 | >50 | mixed | Phased mosaic diploid |
| GRCh38 (2015) | 156 | >50 | mixed | Phased mosaic diploid |
| GRCh38 (2017) | 161 | >50 | mixed | Phased mosaic diploid |

The different numbers of intact LINE-1s in different assemblies could reflect true diversity amongst these individuals, but technical errors in assembly also likely contribute to this variation. Although the CHM genomes improve the assembly of transposable elements, particularly those nested in other repeats (Chaisson et al. 2015), assembly uncertainties or errors around LINE-1 insertion sites in any of these assemblies could underlie the differences in the LINE-1s between assemblies. To investigate, we assessed the integration sites of LINE-1s that are present in the CHMs but absent in GRCh38. We found that none of these integration neighborhoods fall at scaffold ends or major gaps of the GRCh38 assembly. Further, although the CHM genomes contain a similar number of intact LINE-1s as GRCh38, 27 of the CHM1 and 28 of the CHM13 intact LINE-1s correspond to an empty site without a LINE-1 insertion in the reference genome (Table S1). Presumably, some of this variation derives from the uniqueness of the CHM donors relative to the individuals used in the ‘mosaic haploid’ assembly of GRCh38. In order to portray variation amongst individuals, GRCh38 incorporates alternate alleles for regions of the genome that are polymorphic amongst individuals. Indeed, 13 of the intact LINE-1s from CHM1 also present in GRCh38 were found on the ‘alternative’ contigs (representing alternative haplotypes of very polymorphic regions). The corresponding chromosomal assembly these ‘ALT’ contigs does not contain the LINE-1 insertion, reflecting the polymorphic state of these insertions.

We also compared our non-reference intact LINE-1s to the set of intact LINE-1s from a published approach that found potentially active polymorphic LINE-1s based on screening of fosmid libraries of various populations (Beck et al. 2010). From these polymorphic sequences, we found 6 intact LINE-1s that are present in the CHM assemblies but absent from the GRCh38 reference assembly (Table S2). Nonetheless, 37 of the intact LINE-1s in the CHM genomes are not found in GRCh38 or two previous catalogs of intact LINE-1 sequences (Table S2) (Beck et al. 2010; Brouha et al. 2003). This large number of new intact LINE-1s suggests the CHM assemblies provide a major gain in resolution of the intact LINE-1 load in human genomes.

### An estimation of the number of intact LINE-1s in a diploid human genome

While a comparison of intact LINE-1s amongst assemblies supports our use of long-read sequencing of nearly homozygous genomes and provides an indication of population diversity of LINE-1s, the fundamental differences between these assemblies and others prevents a fair comparison of their ability to resolve LINE-1s (Table 1). However, a comparison of CHM1 and CHM13, assemblies generated in very similar manners, should enable a first order guess at the content and variation of LINE-1 that could exist within a heterozygous diploid genome. With 148 and 142 intact LINE-1s in CHM1 and CHM13, respectively, an additive approach to estimating the total number of intact LINE-1s in a simulated diploid of these two genomes gives a total of 290 intact LINE-1 ‘alleles’. We assigned allelic pairs of LINE-1s between CHM1 and CHM13 based upon synteny of each sequence and found that these 290 intact LINE-1 alleles reside at 194 loci in these genomes. Next, we compared the status of each intact LINE-1 locus between the two genomes: intact in both genomes, present in both genomes but only intact in one (sequence polymorphisms), or only present in one genome (insertion polymorphisms). Most of the intact LINE-1s in the CHM genomes have an allelic counterpart in the other CHM genome (126/148 in CHM1 and 117/142 in CHM13; Figure 3A). Of these shared LINE-1s, 96 are intact in both CHM1 and CHM13 (Figure 3A, overlap of dark blue and dark orange; Figure 4, thick lines joining tips). Another 30 loci with intact LINE-1s in CHM1 also contain a LINE-1 in CHM13, but the LINE-1s at these loci in CHM13 have accumulated inactivating mutations (polymorphic in their intactness); CHM13 contains 21 intact LINE-1s that are present but not intact in CHM1 (Figure 3A, “not intact”). Finally, there are 25 loci in CHM1 and 11 loci in CHM13 that contain unique and likely new LINE-1 insertions. These ‘insertionally polymorphic’ loci contain an intact LINE-1 in one genome but are empty in the other genome (Figure 3A, “absent”; Figure 4, blue circles).

**Figure 3.**
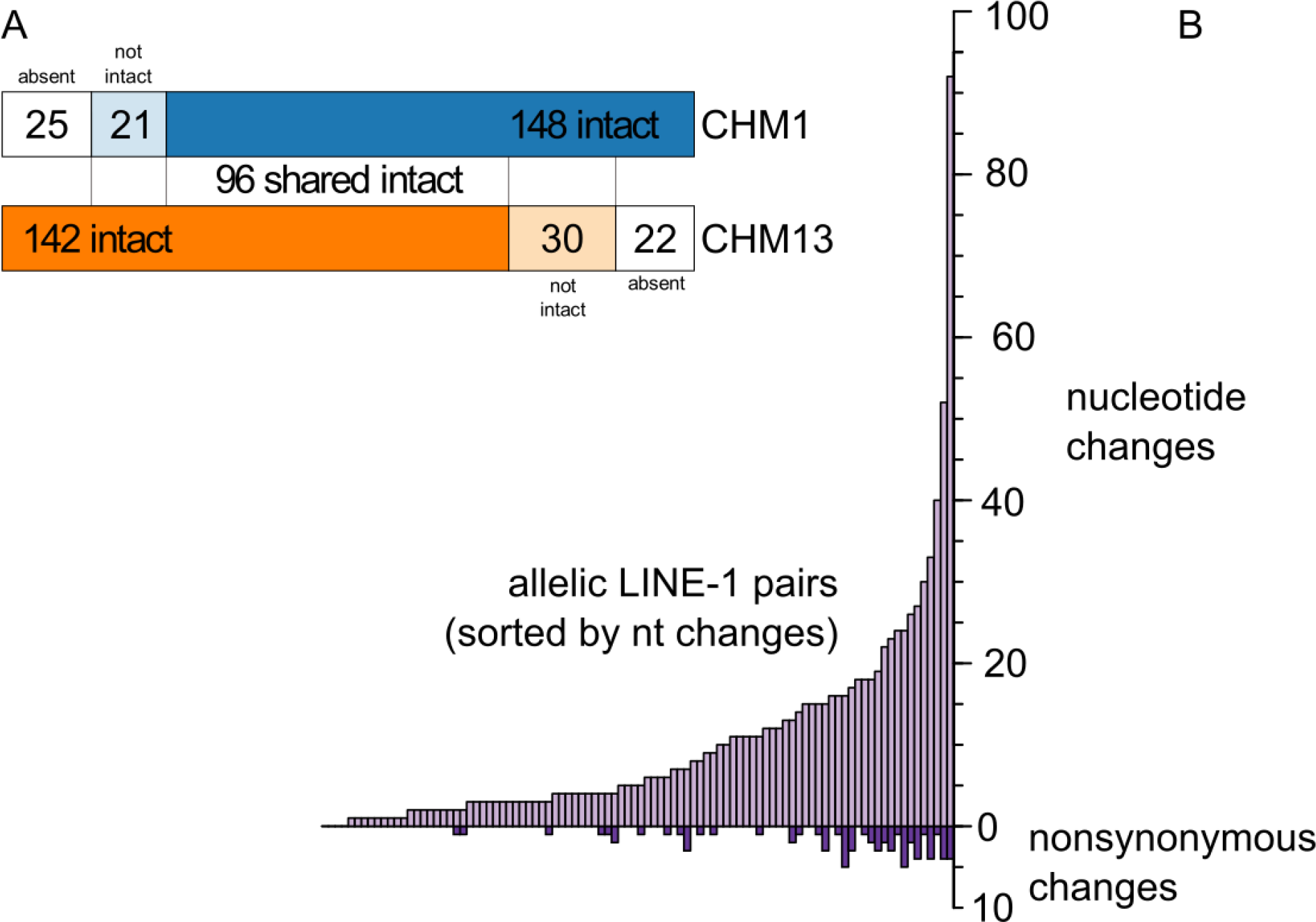
A comparison of intact allelic LINE-1 pairs across CHM genome assemblies. **A.** A liftover comparison of the intact LINE-1s in two nearly homozygous genomes (CHM1, top/blue, and CHM13, bottom/orange) shows 96 intact LINE-1s are shared. Of the LINE-1s that are intact in only one genome, some are present but not intact in the other genome (lighter shading), and some are not present in the other genome (white boxes). **B.** Distribution of nucleotide (top) and non-synonymous (bottom) changes between CHM1 and CHM13 of the 96 allelic intact LINE-1s.

**Figure 4.**
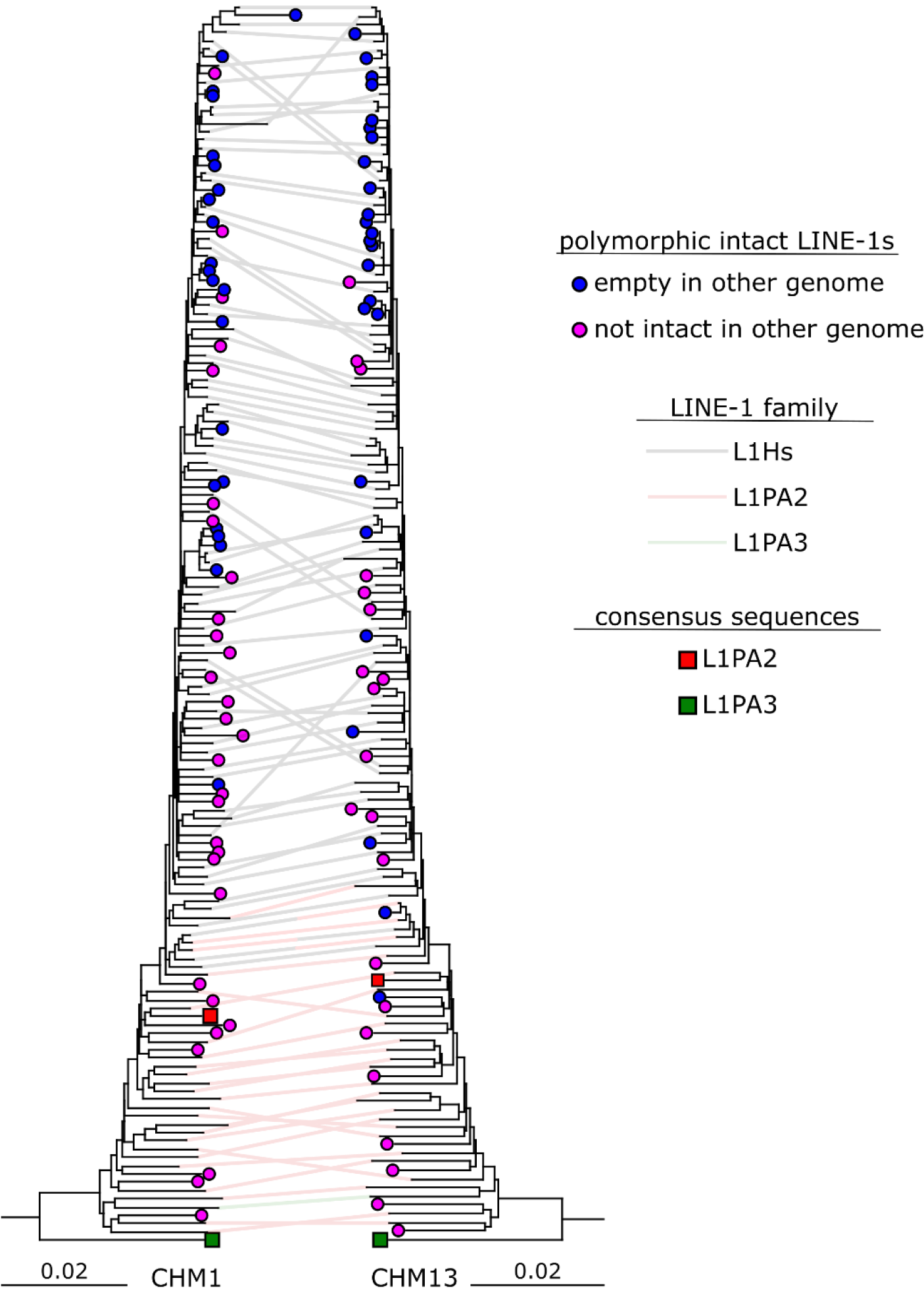
Allelic pairing, new insertions, and phylogeny of intact LINE-1s in two homozygous genomes. A maximum likelihood tree of the complete nucleotide sequence of intact LINE-1 sequences from CHM1 (left) or CHM13 (right) was rooted on a consensus sequence of an older LINE-1 subfamily (L1PA3). Generally, older elements are at the bottom of the tree, and the youngest elements at the top. Elements that are intact in only one genome are marked with a filled circle, where pink corresponds to elements that are present but not intact in the other genome, and blue shows elements that are new insertions relative to the other genome. When a LINE-1 is intact in both genomes, a line connects allelic copies. For shared intact copies, the color of the joining line depicts the LINE-1 family as called by RepeatMasker.

In addition to LINE-1s that are polymorphic in their intactness or presence, we were also able to resolve allelic differences within those loci that contain intact LINE-1s in both CHM1 and CHM13. Young LINE-1s are highly polymorphic in their presence in the human population (Stewart et al. 2011; Beck et al. 2010; Sudmant et al. 2015; Kidd et al. 2010), However, little is known about sequence-level variation in shared LINE-1 insertions. Some diversity of LINE-1 alleles at a single locus has been described (Lutz et al. 2003), but the CHM assemblies provide an unprecedented dataset to analyze this form of diversity across all shared LINE-1s using a simulated pseudodiploid genome. We compared the sequence of pairs of allelic intact LINE-1s in CHM1 and CHM13. The 96 intact LINE-1 allelic pairs show up to five amino acid differences and up to 92 nucleotide differences (Figure 3B). Some of the most different allelic pairs at the nucleotide level have large deletions in the UTRs, which are counted as multiple nucleotide differences in the alignment, but likely represent single deletion events.

Others have deletions and SNPs distributed throughout the UTRs and coding sequence. Only 4/96 allelic pairs of LINE-1s are identical at the nucleotide level; 62/96 pairs are identical at the amino acid level. Overall, allelic differences are pervasive amongst the allelic LINE-1s in the two CHM genomes.

### Evolution of LINE-1s from CHM1 and CHM13

Using the sequences of all intact LINE-1s from each CHM assembly, we generated phylogenetic trees for each genome’s LINE-1s. The resulting trees, rooted with the consensus sequence of an old LINE-1 subfamily L1PA3, largely reflects previous studies and subfamily designations of each sequence from RepeatMasker (Figure 4, black, red, and green lines). Those elements in the oldest part of the trees are also those identified as L1PA2 and L1PA3 (5.6-15.8 million years old) (Khan et al. 2006) while the youngest elements are those identified as L1HS (HS for human-specific) in RepeatMasker. However, the phylogenetic reconstructions show a few areas of disagreement with established subfamily designations. For example, in both trees there is a lack of monophyly for L1HS, with older L1HS and younger L1PA2 falling within the same clade. Additionally, some elements identified as L1PA2 in CHM1 are allelic with elements identified as L1HS in CHM13 and vice versa, likely reflecting absence or mutation of subfamily-classifying nucleotides in some of these sequences. Taken together, these data support a gradual shift from L1PA2 to L1HS.

We also analyzed the distribution of allelic LINE-1s in CHM1 and CHM13 on the two phylogenies (Figure 4, allelic tips joined by a line). With more time elapsed, there is a higher chance that a LINE-1 acquires an inactivating mutation. It follows that we see elements in the older portions of the tree are likely to be shared but are more likely to be polymorphic in their intactness (Figure 4, enrichment of magenta circles in bottom of tree). In contrast, elements in the younger portion of the trees are more likely to be insertionally polymorphic between the two genomes (Figure 4, enrichment of blue circles in the top of tree).

## Discussion

Transposable elements are a major contributor to genetic variation in human genomes (Beck et al. 2011; Auton et al. 2015). As the only highly active and autonomous element in human genomes, LINE-1s impact the host genome through a variety of mechanisms including insertional mutation (Hancks and Kazazian 2012), spreading of epigenetic marks (Grandi et al. 2015), new regulatory sequences (Jacques et al. 2013), serving as a substrate of long non-coding RNAs (Kapusta et al. 2013), and mobilizing other transposable elements (Dewannieux et al. 2003). Because these mechanisms depend on the inserted LINE-1 sequence and/or the precise insertion site, the sequence, location, and variation of the potentially ‘active’ LINE-1s are all crucial factors for understanding the genomic impacts of LINE-1. Previously, tremendous effort has been spent to investigate potentially active LINE-1s in the human genome and polymorphic insertions amongst genomes. However, these studies were plagued by the fundamental shortcomings of an incompletely sequenced human genome, lacking both resolution of repeat regions and phasing of alleles. Here, we capitalize on published deep, long-read assemblies of a nearly homozygous genome to generate a nearly comprehensive catalog of the LINE-1 content of the human genome.

Previously, approaches based on PCR-amplification of the junction of LINE-1s and their integration site, such as junction fragment sequencing (Iskow et al. 2010) and DNA microarrays (Huang et al. 2010), were used to investigate LINE-1 insertion site polymorphism in the human population. However, these approaches only resolve the location and ends of a LINE-1 sequence. Fosmid-based approaches provided a wealth of full-length polymorphic LINE-1 sequences (Beck et al. 2010; Kidd et al. 2010) but lack resolution of allelic variation in the complete set of fixed and polymorphic intact LINE-1s in any single genome. Assembly-free approaches have been used to study the evolution of LINE-1s (Gu et al. 2008; Yang et al. 2014; Platt et al. 2018; de Koning et al. 2011; Smit et al.), which are usually based on the count and identify of short reads or k-mers to consensus or reconstructed ancestral LINE-1 lineages. However, LINE-1 insertion sites and haplotypes remained elusive with these approaches due to the short nature of the input sequence reads. Some LINE-1 data based on various versions of the reference genome are also available (Ewing and Kazazian 2011; Beck et al. 2010; Ewing and Kazazian 2010), but a reference genome does not represent any individual due to the usage of a cohort of donors and other caveats of assembly methods for reference assemblies.

Overall, to our knowledge, this study is the first curation of the collection of intact LINE-1s, including their sequence and location, in two haploid human genomes. Our collection of LINE-1s overlaps with previous efforts to find structural variants in the CHM genomes (Huddleston et al. 2017; Chaisson et al. 2015), and all the sequences we catalog have been previously reported as part of the CHM assemblies. However, the curation of these sequences, not just as RepeatMasker LINE-1 calls, but with an in-depth analysis of sequence characteristics required for LINE-1 retrotransposition, provides the first step towards understanding the functional consequences of the substantial sequence variation in the LINE-1s of individuals.

Our count of 290 intact LINE-1s represents our best guess at the true number of intact LINE-1s in a diploid human genome. However, the actual number is likely even higher. Although we were able to retrieve additional intact LINE-1s that are absent in the reference assembly, including several in the centromeric or simple repeat-containing regions of the genome, there could still be additional LINE-1s present in the haploid CHM genomes. The CHM assemblies include curated BAC sequences spanning segmental duplications that make them especially suited to our analysis over traditional *de novo* assemblies of long reads. There remain, however, regions of the genome refractory to these sequencing and assembly methods which likely contain intact LINE-1s, including long segmental duplications, deep centromeric repeats, and constitutive heterochromatic regions. It is also likely that we miss some intact LINE-1s due to sequencing errors. In particular, PacBio tends to accumulate indels in homopolymer tracts (Hebert et al. 2018), a sequence pattern present at several locations in the LINE-1 sequence. Ultimately, the final count of intact LINE-1s in a single genome awaits a fully phased telomere-to-telomere assembly of an individual human genome.

With the catalog of the intact LINE-1s of two haploid genomes, we combined these two genomes to infer the landscape of intact LINE-1s in a diploid genome. This pseudodiploid of CHM1 and CHM13 contains 290 intact LINE-1 alleles at 194 loci, with 96 loci where both alleles contain intact LINE-1s and 25 or 22 loci with private intact LINE-1s. These numbers update both the total number of LINE-1s in the human genome and the potential level of polymorphism. Though the number of intact LINE-1s from the CHM genomes is not directly comparable to those obtained by the other approaches in Table 1, we highlight several sources that likely contribute to the observed differences. Assemblies based on diploid samples are prone to errors and uncertainty when allelic variation exists. This problem is more pronounced with insertionally polymorphic transposable elements - the allele without the insertion is likely preferred due to the greedy nature of most assembly algorithms. Consistent with this assumption, we observe that the number of intact LINE-1s in diploid-based assemblies (69 and 84, Table 1) is on par with the number of homozygous intact LINE-1s in our pseudodiploid (96, Figure 3); this could be explained by a failure of the assemblies of diploid genomes to resolve heterozygous insertions. Further, given the many individual genomes that comprise the ‘mosaic haploid’ human reference assembly, it likely reflects an ‘average’ haploid version of a genome but is not identical to any single haploid genome. Moreover, the increased number of intact LINE-1s in more recent versions of the reference genome likely reflects increased completeness from advancements in sequencing and assembly technologies.

To identify candidate retrotransposition-competent LINE-1s, we defined ‘intact’ LINE-1s based on the length distribution of intact LINE-1 ORFs. While we include several sequences that have longer or shorter ORFs that still align to the entire length of our reference ORFs (Figure 2C, magenta-outlined pixels), most of these sequences are singletons, suggesting they have not replicated. The only other density of sequences we observe are within one codon of full-length ORFs. Among LINE-1s we classify as non-intact, we do observe multiple sequences with ORF1s 90 or 120 codons shorter than L1_RP_ and ORF2s that are 98 or 265 codons shorter than L1_RP_ ORF2 (Figure S1), representing sequences that have lost a start codon and now have a putative ORF called from the second methionine in the sequence. There could be in-frame indels in the protein-coding sequences of a LINE-1 that do not inactivate retrotransposition, but we observe very little of this variation (135 of the intact LINE-1s have the exact same ORF lengths). The rapid decay away from the ORF lengths of 338 and 1275 codons (evident in the sparseness of the ORF length matrix) is striking and may reflect strong selection for ORFs of these lengths.

The presence of two intact ORFs is only one of the necessary conditions for the *in vivo* retrotransposition of LINE-1. Other conditions include an active promoter in the 5’UTR of LINE-1, absence of more subtle missense debilitating mutations in seemingly intact ORFs, a favorable genomic context of the LINE-1 insertion, and a lack of host restriction mechanisms, amongst others. Indicative of a major transition in LINE-1 evolution, we noticed that the LINE-1 length distribution has two peaks around 6,100 bp that are approximately 130 bp apart (Figure 2B). These peaks correspond to a deletion in the 5’UTR that occurred at the evolutionary transition between the L1PA3 and L1PA4 subfamilies that significantly affects retrotransposition rate, presumably due to the binding of a host restriction factor (ZNF93) to the deleted region (Jacobs et al. 2014). Within our intact LINE-1s, we observe other forms of variation in the 5’ UTR, which certainly could render these elements inactive, via host restriction or loss of regulatory sequences. Clearly, the LINE-1s we classify as intact will not all be competent for retrotransposition. However, the retrotransposition-competent LINE-1s, or the LINE-1s that are replicating in humans now and in the near future, should be among our intact collection. Previous work which cloned many LINE-1 sequences based upon an early version of the human reference assembly found around 50% of intact LINE-1s were active in an *in vitro* assay for retrotransposition. While these studies used a different definition of intactness (>98% sequence identity to L1.3), extrapolating this finding to our dataset would predict around 145 LINE-1 alleles in our pseudodiploid would have measurable *in vitro* activity.

Between the two haploid genomes, we found pervasive variation in the sequence of allelic LINE-1s. Some alleles differ in large deletions at one allele, including deletions of 80bp, 15bp, and 11bp in the 5’ UTRs of one of the allelic pairs. Other pairs differ by as many as 33 nucleotide changes scattered throughout the LINE-1 sequence with no major indels. While allelic diversity has not previously been described widely for LINE-1s, previous literature shows that different common LINE-1 alleles from the same locus can exhibit up to 16-fold differences in retrotransposition assays (Lutz et al. 2003). Further, the repertoire of LINE-1 alleles in any individual can differ dramatically in their cumulative retrotransposition rate (Seleme et al. 2006). Our data also suggests that 32-35% of the intact LINE-1s in one CHM genome are not intact in the other, including the youngest group within L1HS which contains the previously known ‘hot’ LINE-1s (Brouha et al. 2003). Therefore, the collection of retrotransposition-competent LINE-1s in one genome could be very different from those in another genome.

The intact LINE-1s in the two nearly homozygous genomes reflect the retrotransposition history of LINE-1s since the divergence of the two donor individuals. Most of the insertionally polymorphic intact LINE-1s belong to the youngest sub-lineages of L1HS, suggesting that the youngest LINE-1s are responsible for the majority of retrotransposition. Conversely, most of LINE-1s insertions that only differ in their intactness belong to L1PA2 and the older sub-lineages of L1HS, suggesting that, as expected, LINE-1s lose their intact ORFs in some time-dependent fashion. Indeed, by looking at the LINE-1 sequences greater than 5,000 bp in the CHM1 genome, 112 of the 357 L1HS sequences are intact (~31.4%), whereas only 35 of the 1,090 L1PA2 sequences (~3.2%) are intact.

The comparison between phylogenies of intact LINE-1s in CHM1 and CHM13 allows us to visualize interesting patterns in the activity and evolution of LINE-1 in these genomes. Overall, allelic L1s have evolved similarly in the two genomes, evidenced by the similar topology of the two trees and the small number of crossing orthology-indicating lines (Figure 4). The few crossing lines we do see could result from the way we display the tree as a ladder, phylogenetic uncertainty, convergent evolution, or recombination amongst LINE-1s. Perhaps most interestingly, we observe several clusters of LINE-1s that are unique to one genome (insertionally polymorphic, Figure 4, blue circles). These related and recently active LINE-1s could share a highly active parent element or may have sequence changes that make these elements particularly active. While the ethnic origin of CHM13 was not released, previous analysis suggests that CHM1 and CHM13 are closely related. As such, this analysis reflects a small portion of the variation in LINE-1s amongst diverse humans. Indeed, a recent preprint analyzing phased assemblies of three trios of individuals finds an average of 190 intact LINE-1s in individuals of diverse ethnic backgrounds, with only 56 intact LINE-1s shared amongst all three genomes (Chaisson et al. 2018).

In generating an accurate description of even the repeat-rich portions of the human genome, the sequencing and assembly methods used to generate the CHM genomes also provide the data to finally characterize, at allelic resolution, the sequence of major transposable elements like LINE-1 in the human genome, a key step in understanding the sequence and functional diversity of LINE-1s in human biology. This catalog of LINE-1 sequences provides the foundation for countless future studies of the evolution and functional variation of LINE-1s in recent human evolutionary history.

## Methods

### Retrieving LINE-1 from the haploid and reference genomes

LINE-1s were identified according to the RepeatMasker annotation from CHM1 and CHM13 assemblies (GenBank assembly accession: GCA_001297185.2 and GCA_000983455.2). BLAST searches of CHM1 and CHM13 used L1.3 (GenBank accession number: L19088) as a query. The RepeatMasker annotation was filtered for keyword “L1’ in the “matching repeat” column and converted to bed format. Subsequently, LINE-1 sequences were retrieved from the genome according to the bed file using the “subseq” function of seqtk (https://github.com/lh3/seqtk). GRCh37 and GRCh38 reference genome sequences and RepeatMasker annotations were downloaded from the annotations of GenBank assembly GCA_001297185.2 and GCA_000983455.2 and processed in a similar manner as the CHM genomes to find LINE-1s. LINE-1s on the ALT contigs of the reference genomes were manually inspected for their corresponding chromosomal location if possible and assigned as an alternative LINE-1 allele of that chromosomal location in the reference genome.

### Intact LINE-1 identification

Full-length LINE-1s (Files S1 and S2) were found by filtering the RepeatMasker annotation of the CHM genomes requiring the length of the annotated LINE-1 sequence to be equal to or longer than 5,000 bp. LINE-1 ORFs were found by using EMBOSS (Rice et al. 2000) “getorf” function on full-length LINE-1s with “-find 1” setting to return the translated sequences of the ORFs. The translated ORFs were subsequently searched using BLASTp with the translated ORFs of L1_RP_ (GenBank accession number: AF148856). For each of the full-length CHM1 and CHM13 LINE-1, a custom perl script processed the BLASTp output to find the ORF of the LINE-1 that forms the longest alignment to the ORF1 and ORF2 protein of L1_RP_. LINE-1s with intact ORFs were identified in the distribution of the longest called ORFs of each LINE-1 that align to the reference ORFs, which correspond to ORF1 length of 338 codons and ORF2 length of 1,275 codons. Singletons near these ORF lengths were manually inspected to find additional ORFs that align to the full length of the L1_RP_ reference ORFs.

### Synteny of LINE-1 in haploid and reference genomes

CHM1 and CHM13 genome sequences were aligned, chromosome by chromosome, to the GRCh37 and GRCh38 reference genomes using lastz (Harris 2007) under the setting of “--notransition --step=20 --format=lav”. The alignments were then processed using lavToPsl, axtChain, and chainMergeSort functions of the UCSC genome browser utilities (http://hgdownload.soe.ucsc.edu/admin/exe/) with default settings. LINE-1 coordinates on CHM1 and CHM13 were subsequently converted to GRCh37 and GRCh38 coordinates with the default setting of the liftOver tool of the UCSC genome browser utilities and the processed chain file mentioned above. For the LINE-1s that could not be directly lifted-over with the default liftOver setting, we took the sequences that are 2,000 bp from each end of LINE-1s and lifted them over to the reference genomes. For this step, the lifted-over coordinates of both extended ends were in the same neighborhood of the target reference genome for almost all LINE-1s. We were unable to obtain coordinates for a small subset of LINE-1s flanked by repetitive sequence using this method, and so are unable to assign a genomic position and to pair possible allelic variants; one exceptional LINE-1 flanked by a repeat-rich region had enough unique sequence on both sides to assign as allelic in the CHM1 and CHM13 genomes but still could not be assigned to a genomic coordinate (Table S2, row 236).

### Phylogeny of LINE-1s

Intact LINE-1s from CHM1 and CHM13 were aligned using MUSCLE (Edgar 2004) with default settings. Based on our ORF analyses, 148 intact CHM1 LINE-1s and 142 intact CHM13 LINE-1s were input for alignment and phylogenetic reconstruction. A model test was performed through R using the phangorn package (Schliep 2011) and the GTR model was chosen using AIC criteria. Maximum likelihood phylogenies were generated using the phangorn package of R. First, a Neighbor Joining tree was generated and used as the starting tree for the ML analysis, then ML analysis was performed. After ML analysis, 100 bootstrap replicates were performed.

## Supplementary Materials

Figure S1

Heatmap of translated ORF lengths in full-length LINE-1s from CHM1.

A zoomed-out version of Figure 1C shows the distribution of all called translated ORF lengths in the complete set of full-length LINE-1 sequences (>5000bp). Many full-length sequences have either an intact ORF1 or ORF2, but only 148 have both intact ORFs that align along the entire length of a reference amino acid sequence (L1_RP_).

Figure S2

Identification of intact LINE-1s in CHM13.

**A.** The distribution of lengths of LINE-1-masked sequences shows most sequences are less than 1,000 bp. An additional peak around 6,000 bp contains the full-length LINE-1 copies. **B.** A zoomed-in view of the boxed region of the complete length histogram. Intact LINE-1 sequences are highlighted in magenta. **C.** A heatmap of the number of sequences with the indicated translated ORF1 and ORF2 lengths. Within the full-length set of LINE-1 sequences, only 142 sequences (magenta-outlined pixels) encode putative ORFs that align along the entire length of the ORF1p and ORF2p sequences of a reference element (L1_RP_).

Table S1

A comparison of intact LINE-1s across GRCh38 and CHM genome assemblies. Counts of the number of LINE-1s that are intact, present but not intact (non-intact), and absent are shown. Only LINE-1s that are intact in one of the genomes in each comparison are counted - ‘NA's represent the categories where LINE-1s are intact in none of the genomes in each comparison.

Table S2

Analysis of intact LINE-1 alleles from GRCh38 and CHM genome assemblies. Coordinates of LINE-1s that are intact in either CHM1, CHM13, or GRCh38 (hg38) are shown. GRCh37 (hg19) coordinates are based on liftOver from GRCh38 coordinates. ‘Intact’ represents the LINE-1s with intact ORFs, ‘short’ represents the LINE-1s that are present at the insertion site but the ORFs are no longer intact, and ‘.’ indicates that the site does not contain the LINE-1 insertion.

File S1

Nucleotide sequence alignment of CHM1 intact LINE-1s and representative L1PA2-L1PA6 as outgroup

File S2

Nucleotide sequence alignment of CHM13 intact LINE-1s and representative L1PA2-L1PA6 as outgroup

File S3

Maximum likelihood phylogeny based on the nucleotide sequence alignment of CHM1 intact LINE-1s and representative L1PA2-L1PA6 as outgroup

File S4

Maximum likelihood phylogeny based on the nucleotide sequence alignment of CHM13 intact LINE-1s and representative L1PA2-L1PA6 as outgroup

## Acknowledgments

We thank John Huddleston, Michael Metzger and members of the Metzger lab, members of the McLaughlin lab, and Janet Young for technical advice and critical comments on this manuscript.

## Funding

This work was supported by a grant from the National Institutes of Health (NIGMS R00 GM112941 to RNM).

## Disclosure Declaration

The authors have nothing to declare.

## References

Audano PA, Sulovari A, Graves-Lindsay TA, Cantsilieris S, Sorensen M, Welch AE, Dougherty ML, Nelson BJ, Shah A, Dutcher SK, et al. 2019. Characterizing the Major Structural Variant Alleles of the Human Genome. Cell 176: 663–675.e19.

Auton A, Abecasis GR, Altshuler DM, Durbin RM, Bentley DR, Chakravarti A, Clark AG, Donnelly P, Eichler EE, Flicek P, et al. 2015. A global reference for human genetic variation. Nature 526: 68–74.

Beck CR, Collier P, Macfarlane C, Malig M, Kidd JM, Eichler EE, Badge RM, Moran J V. 2010. LINE-1 Retrotransposition Activity in Human Genomes. Cell 141: 1159–1170.

Beck CR, Garcia-Perez JL, Badge RM, Moran J V. 2011. LINE-1 Elements in Structural Variation and Disease. Annu Rev Genomics Hum Genet 12: 187–215.

Bejerano G, Lowe CB, Ahituv N, King B, Siepel A, Salama SR, Rubin EM, Kent WJ, Haussler D. 2006. A distal enhancer and an ultraconserved exon are derived from a novel retroposon. Nature 441: 87–90.

Brouha B, Schustak J, Badge RM, Lutz-Prigge S, Farley AH, Moran J V., Kazazian HH. 2003. Hot L1s account for the bulk of retrotransposition in the human population. Proc Natl Acad Sci.

Canapa A, Barucca M, Biscotti MA, Forconi M, Olmo E. 2015. Transposons, Genome Size, and Evolutionary Insights in Animals. Cytogenet Genome Res 147: 217–39.

Chaisson MJP, Huddleston J, Dennis MY, Sudmant PH, Malig M, Hormozdiari F, Antonacci F, Surti U, Sandstrom R, Boitano M, et al. 2015. Resolving the complexity of the human genome using single-molecule sequencing. Nature 517: 608–611.

Chaisson MJP, Sanders AD, Zhao X, Malhotra A, Porubsky D, Rausch T, Gardner EJ, Rodriguez O, Guo L, Collins RL, et al. 2018. Multi-platform discovery of haplotype-resolved structural variation in human genomes. bioRxiv 193144.

Chow JC, Ciaudo C, Fazzari MJ, Mise N, Servant N, Glass JL, Attreed M, Avner P, Wutz A, Barillot E, et al. 2010. LINE-1 activity in facultative heterochromatin formation during X chromosome inactivation. Cell 141: 956–69.

Crow MK. 2010. Long interspersed nuclear elements (LINE-1): potential triggers of systemic autoimmune disease. Autoimmunity 43: 7–16.

de Koning AP, Gu W, Castoe TA, Batzer MA, Pollock DD. 2011. Repetitive elements may comprise over two-thirds of the human genome. PLoS Genet 7: e1002384.

Dewannieux M, Esnault C, Heidmann T. 2003. LINE-mediated retrotransposition of marked Alu sequences. Nat Genet 35: 41–48.

Dombroski BA, Scott AF, Kazazian HH. 1993. Two additional potential retrotransposons isolated from a human L1 subfamily that contains an active retrotransposable element. Proc Natl Acad Sci U S A 90: 6513–7.

Edgar RC. 2004. MUSCLE: Multiple sequence alignment with high accuracy and high throughput. Nucleic Acids Res 32: 1792–1797.

Ewing AD, Kazazian HH. 2010. High-throughput sequencing reveals extensive variation in human-specific L1 content in individual human genomes. Genome Res 20: 1262–1270.

Ewing AD, Kazazian HH. 2011. Whole-genome resequencing allows detection of many rare LINE-1 insertion alleles in humans. Genome Res.

Fan JB, Surti U, Taillon-Miller P, Hsie L, Kennedy GC, Hoffner L, Ryder T, Mutch DG, Kwok PY. 2001. Paternal origins of complete hydatidiform moles proven by whole genome single-nucleotide polymorphism haplotyping. Genomics 79: 58–62.

Finnerty H, Mi S, Veldman GM, McCoy JM, LaVallie E, Edouard P, Tang X-Y, Howes S, Keith JC, Racie L, et al. 2002. Syncytin is a captive retroviral envelope protein involved in human placental morphogenesis. Nature 403: 785–789.

Grandi FC, Rosser JM, Newkirk SJ, Yin J, Jiang X, Xing Z, Whitmore L, Bashir S, Ivics Z, Izsvák Z, et al. 2015. Retrotransposition creates sloping shores: A graded influence of hypomethylated CpG islands on flanking CpG sites. Genome Res 25: 1135–1146.

Gu W, Castoe TA, Hedges DJ, Batzer MA, Pollock DD. 2008. Identification of repeat structure in large genomes using repeat probability clouds. Anal Biochem 380: 77–83.

Hancks DC, Kazazian HH. 2012. Active human retrotransposons: Variation and disease. Curr Opin Genet Dev 22: 191–203.

Harris RS. 2007. Improved pairwise alignment of genomic DNA. The Pennsylvania State University.

Hebert PDN, Braukmann TWA, Prosser SWJ, Ratnasingham S, DeWaard JR, Ivanova N V., Janzen DH, Hallwachs W, Naik S, Sones JE, et al. 2018. A Sequel to Sanger: amplicon sequencing that scales. BMC Genomics 19: 219.

Huang CRL, Schneider AM, Lu Y, Niranjan T, Shen P, Robinson MA, Steranka JP, Valle D, Civin CI, Wang T, et al. 2010. Mobile interspersed repeats are major structural variants in the human genome. Cell 141: 1171–1182.

Huddleston J, Chaisson MJP, Steinberg KM, Warren W, Hoekzema K, Gordon D, Graves-Lindsay TA, Munson KM, Kronenberg ZN, Vives L, et al. 2017. Discovery and genotyping of structural variation from long-read haploid genome sequence data. Genome Res 27: 677–685.

Iskow RC, McCabe MT, Mills RE, Torene S, Pittard WS, Neuwald AF, Van Meir EG, Vertino PM, Devine SE. 2010. Natural Mutagenesis of Human Genomes by Endogenous Retrotransposons. Cell 141: 1253–1261.

Jacobs FM, Greenberg D, Nguyen N, Haeussler M, Ewing AD, Katzman S, Paten B, Salama SR, Haussler D. 2014. An evolutionary arms race between KRAB zinc-finger genes ZNF91/93 and SVA/L1 retrotransposons. Nature 516: 242–245.

Jacques P-É, Jeyakani J, Bourque G. 2013. The majority of primate-specific regulatory sequences are derived from transposable elements. PLoS Genet 9: e1003504.

Kapusta A, Kronenberg Z, Lynch VJ, Zhuo X, Ramsay LA, Bourque G, Yandell M, Feschotte C. 2013. Transposable Elements Are Major Contributors to the Origin, Diversification, and Regulation of Vertebrate Long Noncoding RNAs ed. H.E. Hoekstra. PLoS Genet 9: e1003470.

Khan H, Smit A, Boissinot S. 2006. Molecular evolution and tempo of amplification of human LINE-1 retrotransposons since the origin of primates. Genome Res 16: 78–87.

Kidd JM, Graves T, Newman TL, Fulton R, Hayden HS, Malig M, Kallicki J, Kaul R, Wilson RK, Eichler EE. 2010. A human genome structural variation sequencing resource reveals insights into mutational mechanisms. Cell 143: 837–47.

Kimberland ML, Divoky V, Prchal J, Schwahn U, Berger W, Kazazian HH. 1999. Full-length human L1 insertions retain the capacity for high frequency retrotransposition in cultured cells. Hum Mol Genet 8: 1557–60.

Lander E, Heaford A, Sheridan A, Linton L, Birren B, Subramanian A, Coulson A, Nusbaum C, Zody M, Dunham A, et al. 2001. Initial sequencing and analysis of the human genome. Nature 409: 860–921.

Lutz SM, Vincent BJ, Kazazian HH, Batzer MA, Moran J V. 2003. Allelic Heterogeneity in LINE-1 Retrotransposition Activity. Am J Hum Genet 73: 1431–1437.

Moran J V, Holmes SE, Naas TP, DeBerardinis RJ, Boeke JD, Kazazian HH. 1996. High frequency retrotransposition in cultured mammalian cells. Cell 87: 917–27.

Platt RN, Vandewege MW, Ray DA. 2018. Mammalian transposable elements and their impacts on genome evolution. Chromosom Res 26: 25–43.

Rice P, Longden I, Bleasby A. 2000. EMBOSS: the European Molecular Biology Open Software Suite. Trends Genet 16: 276–7.

Schliep KP. 2011. phangorn: Phylogenetic analysis in R. Bioinformatics 27: 592–593.

Schneider VA, Graves-Lindsay T, Howe K, Bouk N, Chen H-C, Kitts PA, Murphy TD, Pruitt KD, Thibaud-Nissen F, Albracht D, et al. 2017. Evaluation of GRCh38 and de novo haploid genome assemblies demonstrates the enduring quality of the reference assembly. Genome Res 27: 849–864.

Seleme M del C, Vetter MR, Cordaux R, Bastone L, Batzer MA, Kazazian HH. 2006. Extensive individual variation in L1 retrotransposition capability contributes to human genetic diversity. Proc Natl Acad Sci U S A 103: 6611–6.

Smit A, Hubley R, Green P. RepeatMasker Open-4.0.

Smit AFA, Tóth G, Riggs AD, Jurka J. 1995. Ancestral, mammalian-wide subfamilies of LINE-1 repetitive sequences. J Mol Biol 246: 401–417.

Sotero-Caio CG, Platt RN, Suh A, Ray DA. 2017. Evolution and diversity of transposable elements in vertebrate genomes. Genome Biol Evol 9: 161–177.

Steinberg KM, Schneider VA, Graves-Lindsay TA, Fulton RS, Agarwala R, Huddleston J, Shiryev SA, Morgulis A, Surti U, Warren WC, et al. 2014. Single haplotype assembly of the human genome from a hydatidiform mole. Genome Res 24: 2066–2076.

Stewart C, Kural D, Strömberg MP, Walker JA, Konkel MK, Stütz AM, Urban AE, Grubert F, Lam HYK, Lee W-P, et al. 2011. A Comprehensive Map of Mobile Element Insertion Polymorphisms in Humans ed. H.S. Malik. PLoS Genet 7: e1002236.

Sudmant PH, Rausch T, Gardner EJ, Handsaker RE, Abyzov A, Huddleston J, Zhang Y, Ye K, Jun G, Fritz MH-Y, et al. 2015. An integrated map of structural variation in 2,504 human genomes. Nature 526: 75–81.

Szak ST, Pickeral OK, Makalowski W, Boguski MS, Landsman D, Boeke JD. 2002. Molecular archeology of L1 insertions in the human genome. Genome Biol 3: research0052.

Tang Z, Steranka JP, Ma S, Grivainis M, Rodić N, Huang CRL, Shih I-M, Wang T-L, Boeke JD, Fenyö D, et al. 2017. Human transposon insertion profiling: Analysis, visualization and identification of somatic LINE-1 insertions in ovarian cancer. Proc Natl Acad Sci 114: E733–E740.

Xing J, Zhang Y, Han K, Salem AH, Sen SK, Huff CD, Zhou Q, Kirkness EF, Levy S, Batzer MA, et al. 2009. Mobile elements create structural variation: analysis of a complete human genome. Genome Res 19: 1516–26.

Yang L, Brunsfeld J, Scott L, Wichman H. 2014. Reviving the Dead: History and Reactivation of an Extinct L1. PLoS Genet 10: e1004395.

