## Supplementary figures and images for "Characterization of LINE-1 transposons in a human genome at allelic resolution"

### Supplemental Figure S1

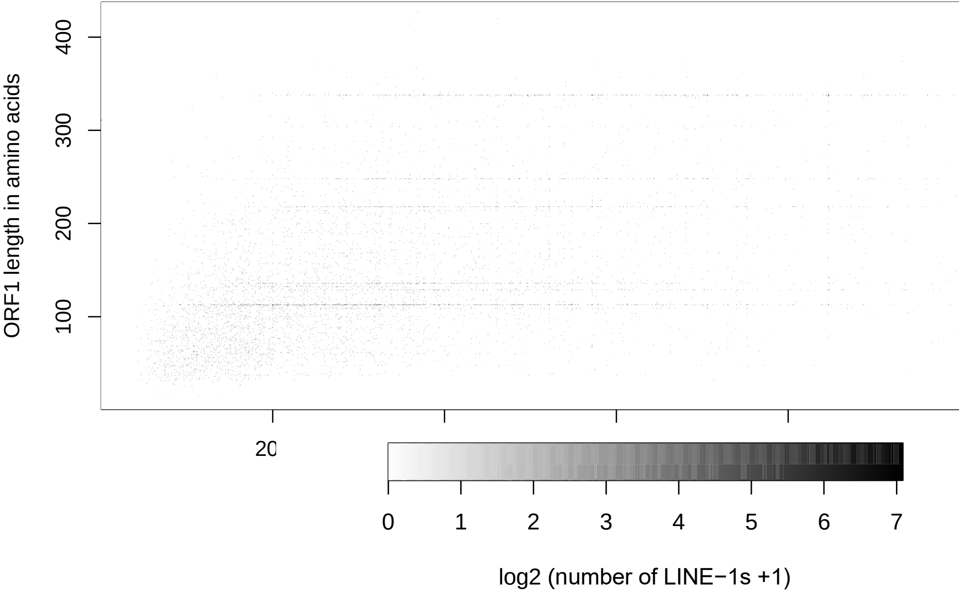

### Supplemental Figure S2

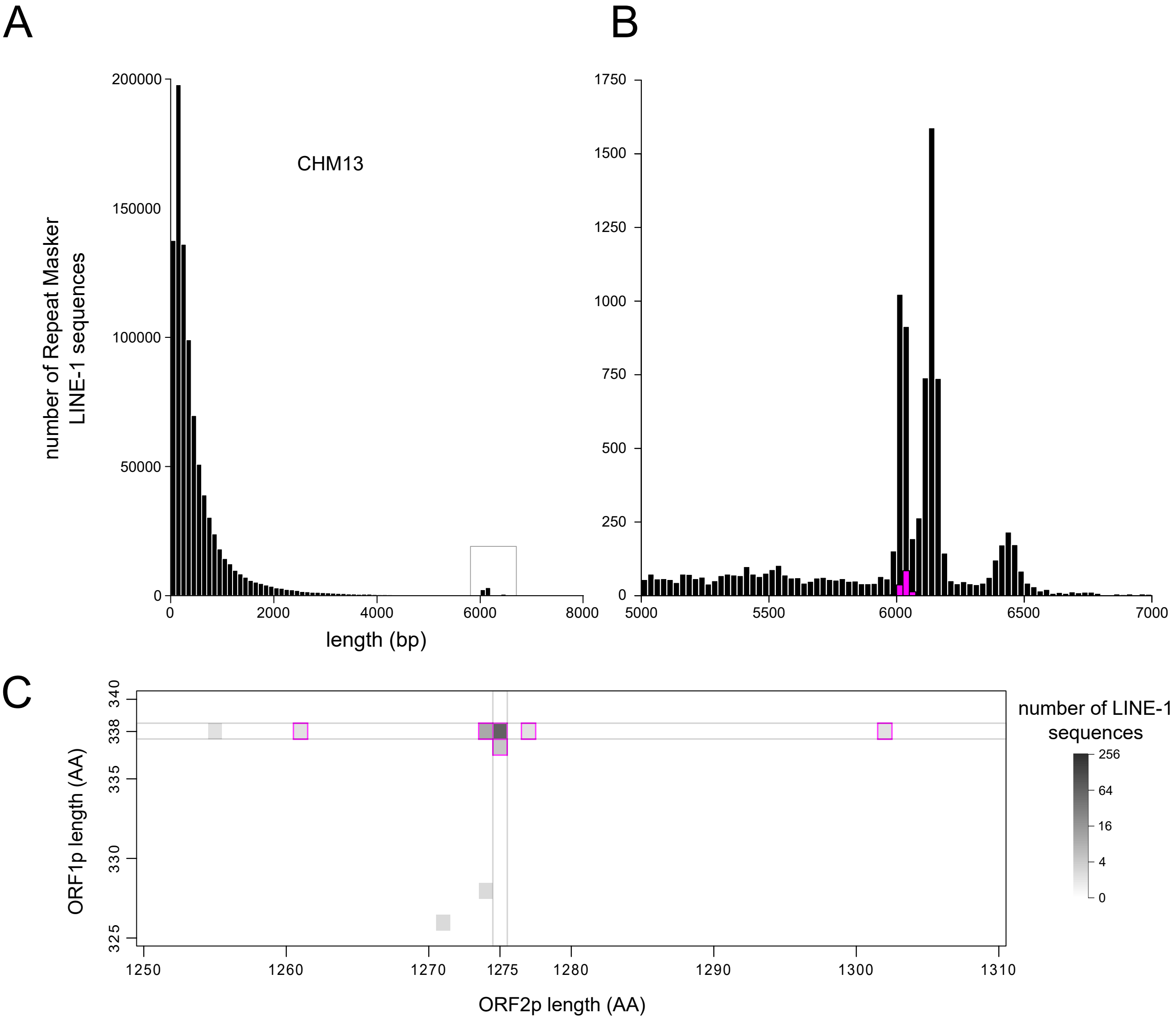
